# A sensory–preoptic circuit drives capsaicin-induced hypothermia

**DOI:** 10.64898/2026.03.21.713372

**Authors:** Hanan Bouaouda, Jan Siemens

## Abstract

Capsaicin, the principal agonist of the heat-sensitive transient receptor potential vanilloid 1 (TRPV1) channel, triggers profound hypothermia by suppressing thermogenesis and promoting heat loss. While TRPV1 has been extensively characterized as a molecular sensor of heat and inflammatory mediators, the neural circuits through which capsaicin drives body cooling remain poorly defined. Emerging work suggests that peripheral TRPV1 activation modulates afferent input to central thermoregulatory networks, yet how —and where— these signals are integrated within the central nervous system to induce hypothermia has remained unresolved. Intriguingly, optogenetic or chemogenetic activation of specific neuronal populations in the preoptic area (POA) of the hypothalamus has been shown to evoke deep hypothermia, raising the possibility that capsaicin engages overlapping thermoregulatory circuits. However, whether capsaicin-driven hypothermia depends on these POA neuron populations has not been directly tested.

Here, we address this question by selectively and permanently silencing defined POA neuronal subtypes and assessing their contribution to capsaicin-evoked hypothermia. Silencing VMPO^LepR^ or POA^Vgat^ neurons attenuated the hypothermic response but did not abolish it, whereas silencing POA^Vglut2^ neurons nearly eliminated capsaicin-induced hypothermia. These findings identify POA^Vglut2^ neurons as a critical central node through which peripheral TRPV1 activation drives body cooling. Together, our results demonstrate that capsaicin induces hypothermia via afferent sensory pathways that converge onto hypothalamic preoptic thermoregulatory neurons, revealing a direct circuit-level link between peripheral TRPV1 signalling and central control of body temperature.

## Introduction

Capsaicin, the pungent ingredient in chili peppers, is a selective agonist of TRPV1 (Jancso-Gabor *et al*., 1970a; Szolcsanyi & Jancso-Gabor, 1975; Caterina *et al*., 1997; Tominaga *et al*., 1998; Szallasi & Blumberg, 1999; Szolcsanyi, 2015). Capsaicin has long been known to trigger hypothermia in mammals. Ever since their studies in the late 1940s and early 1950s, the work by Jancsó-Gabor and colleagues revealed that systemic application of capsaicin to rodents and other small animals caused profound and reversible hypothermia, typically lasting from a few minutes up to several hours (Jancsó, 1955; Jancso-Gabor *et al*., 1970a).

The cloning of TRPV1 has uncovered the molecular mediator of capsaicin’s action on core body temperature (Caterina *et al*., 1997). Targeting TRPV1 with highly selective and efficacious antagonists can have an opposing effect to that observed for capsaicin and can trigger hyperthermia in animal models and humans (Swanson *et al*., 2005; Immke & Gavva, 2006; Gavva *et al*., 2008; Rowbotham *et al*., 2011; Othman *et al*., 2013; Quiding *et al*., 2013; Manitpisitkul *et al*., 2016). TRPV1’s apparent constitutive and tonic role in core body temperature regulation has hampered the entry of TRPV1-selective antagonists into clinical practice for the treatment of pain.

These discoveries prompted a search to identify the anatomical location where TRPV1 regulates core body temperature. While early works speculated that the hypothalamic thermoregulatory centres could be involved in the signalling (Jancso-Gabor *et al*., 1970b; a; Szolcsanyi *et al*., 1971; Nakayama *et al*., 1978; Sasamura *et al*., 1998; Karlsson *et al*., 2005), recent focus has been on its peripheral targeting by the use of restricted TRPV1 modulators (Steiner *et al*., 2007; Tamayo *et al*., 2008). Using conditional, tissue-selective TRPV1 knockout mouse lines, Yue and colleagues showed that TRPV1 receptors localized to peripheral sensory neurons are responsible for bidirectional control of core body temperature elicited by pharmacological agonists and antagonists (Yue *et al*., 2022). These results were subsequently corroborated by a study performing precise peripheral nerve transections that pointed to an abdominal origin of TRPV1-harboring sensory neurons regulating core body temperature (Garami *et al*., 2023). Moreover, both studies suggest that CNS-directed signal transmission of the TRPV1-expressing sensory neurons is required for capsaicin-induced hypothermia, rather than peripheral release of vasoactive (and thereby potentially thermoregulatory) mediators from their nerve endings. Collectively, these findings raise the question of how CNS-directed TRPV1 signals can have such a profound effect on core body temperature and which neuronal circuits are mediating these effects.

Within the CNS, the POA is considered a main hub controlling core body temperature. The overarching neuroanatomical pathways governing afferent information flow to and efferent signals from the POA to modulate the activity of peripheral organ systems to regulate core body temperature are known (Nakamura, 2011; Romanovsky, 2018; Tan & Knight, 2018; Morrison & Nakamura, 2019). However, detailed cellular and circuit-level information, how temperature information is integrated, and how proper outputs are generated to fine-tune core body temperature remain poorly understood.

Several recent studies have implicated a molecularly diverse, partially overlapping neuron population in the ventromedial part of the POA (VMPO) in driving hypothermia in mice (Song *et al*., 2016; Tan *et al*., 2016; Yu *et al*., 2016; Zhao *et al*., 2017; Hrvatin *et al*., 2020; Takahashi *et al*., 2020; Zhang *et al*., 2020). These neurons express the glutamate transporter Vglut2, and a subset of them harbour the leptin receptor LepR. Intriguingly, when activated optogenetically or chemogenetically, these neurons evoke a robust hypothermic response similar to that induced by systemic TRPV1 activation by capsaicin, albeit much more prolonged. Here, we tested the hypothesis that peripheral activation of TRPV1 in sensory neurons mediates hypothermia via signals converging onto hypothalamic POA neurons.

## Results

### VMPO^LepR^ neurons partially mediate the hypothermic effect of capsaicin

In agreement with a large body of literature, acute administration of capsaicin (1 mg/kg, subcutaneous) triggered a robust decrease in core body temperature from 36.06 ± 0.31 °C to 31.61 ± 0.34 °C (Fig. 1A, Table 1), (Jancso-Gabor *et al*., 1970a; Caterina *et al*., 2000; Ayoub *et al*., 2011; Ivanova *et al*., 2023). The hypothermic response peaked at ∼35 min after injection and returned to the baseline level within ∼75 min, whereas vehicle treatment had no effect (Fig. 1A). This hypothermic response was at least partially driven by increased heat dissipation and reduced thermogenesis, as evidenced by elevated tail temperature (cutaneous vasodilation) and decreased brown adipose tissue temperature (Fig. 1B).

**Figure 1.**
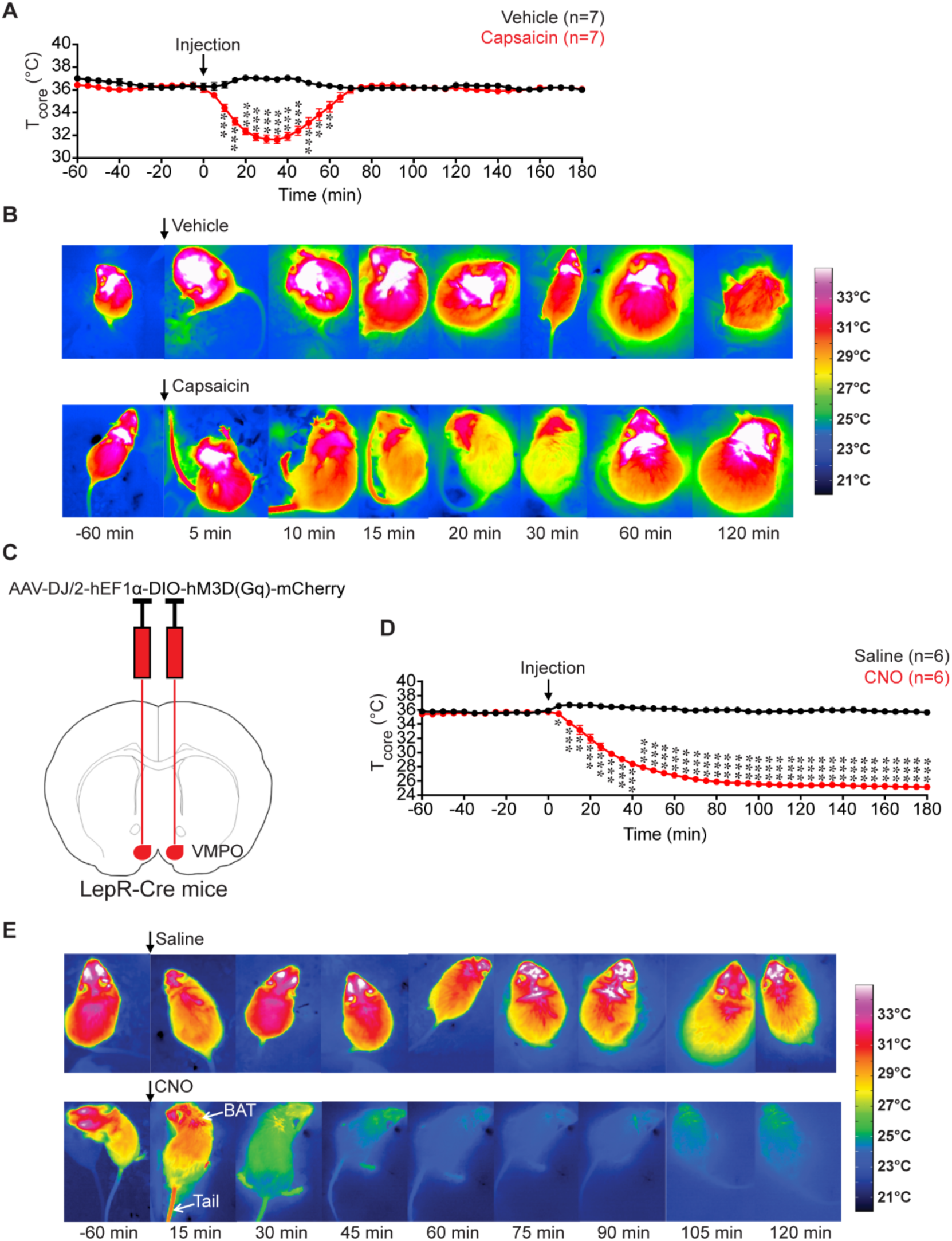
Systemic capsaicin and chemogenetic activation of VMPO^LepR^ neurons induce hypothermia. **(A)** Core body temperature following a single subcutaneous injection of capsaicin (1 mg/kg) or vehicle. Each data point corresponds to the mean value of body temperature ± SEM (n = 7 mice per group). **(B)** Representative infrared thermographic images showing that subcutaneous administration of capsaicin increased tail temperature and decreased BAT temperature 5 min after injection. **(C)** Schematic coronal diagram showing injection of AAV-DJ/2-hEF1α-DIO-hM3D(Gq)-mCherry into the POA of LepR-Cre mice. **(D)** Core body temperature following administration of saline or CNO. Each data point corresponds to the mean value of body temperature ± SEM (n = 6 mice per group). **(E)** Infrared thermography of a LepR-Cre mouse following administration of saline or CNO. Data were recorded every 5 min. Asterisks indicate statistically significant differences (*p < 0.05, **p < 0.01, ***p < 0.001).

**Table1.** Statistical results from repeated measures two-way ANOVA.

|  | Treatment | Time | Interaction |
| --- | --- | --- | --- |
| Fig.1A,<br>T<sub>core</sub> | $F_{1, 12} = 46.58, P < 0.001$ | $F_{4.762, 57.14} = 22.24, P < 0.001$ | $F_{48, 576} = 39.79, P < 0.001$ |
| Fig.1C,<br>T<sub>core</sub> | $F_{1, 10} = 926.1, P < 0.001$ | $F_{2.459, 24.59} = 226.1, P < 0.001$ | $F_{48, 480} = 229.3, P < 0.001$ |
| Fig.2C,<br>T<sub>core</sub> | $F_{1, 12} = 24.23, P < 0.001$ | $F_{4.865, 58.38} = 23.12, P < 0.001$ | $F_{48, 576} = 49.44, P < 0.001$ |
| Fig.2D,<br>T<sub>core</sub> | $F_{1, 12} = 0.4997, P = 0.49$ | $F_{5.406, 64.88} = 6.772, P < 0.001$ | $F_{48, 576} = 9.277, P < 0.001$ |
| Fig.2E,<br>delta<br>T<sub>core</sub> | $F_{1, 12} = 8.246, P = 0.01$ | $F_{4.991, 59.90} = 40.36, P < 0.001$ | $F_{48, 576} = 15.99, P < 0.001$ |
| Fig.3C,<br>T<sub>core</sub> | $F_{1, 12} = 7.441, P = 0.02$ | $F_{5.078, 60.94} = 38.67, P < 0.001$ | $F_{48, 576} = 52.14, P < 0.001$ |
| Fig.3D,<br>T<sub>core</sub> | $F_{1, 12} = 0.2223, P = 0.65$ | $F_{3.832, 45.99} = 4.917, P = 0.002$ | $F_{48, 576} = 1.396, P = 0.04$ |
| Fig.3E,<br>delta<br>T<sub>core</sub> | $F_{1, 12} = 11.51, P = 0.005$ | $F_{4.150, 49.80} = 28.23, P < 0.001$ | $F_{48, 576} = 13.31, P < 0.001$ |
| Fig.4A,<br>Water<br>intake | $F_{1, 7} = 288.8, P < 0.001$ | $F_{2.915, 20.41} = 1.427, P = 0.26$ | $F_{21, 147} = 1.837, P = 0.02$ |
| Fig.4B,<br>Food<br>intake | $F_{1, 7} = 10.34, P = 0.01$ | $F_{2.870, 20.09} = 1.880, P = 0.17$ | $F_{21, 147} = 2.784, P < 0.001$ |
| Fig.4B,<br>Body<br>weight | $F_{1, 7} = 3.094, P = 0.12$ | $F_{1.664, 11.65} = 1.840, P = 0.20$ | $F_{21, 147} = 5.054, P < 0.001$ |
| Fig.5C,<br>T<sub>core</sub> | $F_{1, 4} = 5.527, P = 0.08$ | $F_{1.851, 7.402} = 9.571, P = 0.010$ | $F_{48, 192} = 16.25, P < 0.001$ |
| Fig.5D,<br>T<sub>core</sub> | $F_{1, 6} = 14.36, P = 0.009$ | $F_{3.863, 23.18} = 25.35, P < 0.001$ | $F_{48, 288} = 32.93, P < 0.001$ |
| Fig.5E,<br>delta<br>T<sub>core</sub> | $F_{1, 5} = 1.585, P = 0.26$ | $F_{2.897, 14.49} = 50.14, P < 0.001$ | $F_{48, 240} = 1.728, P = 0.004$ |
| Fig.6C,<br>T<sub>core</sub> | $F_{1, 12} = 11.24, P = 0.006$ | $F_{4.004, 48.05} = 22.03, P < 0.001$ | $F_{48, 576} = 48.56, P < 0.001$ |
| Fig.6D,<br>T<sub>core</sub> | $F_{1, 12} = 1.400, P = 0.26$ | $F_{4.627, 55.52} = 8.975, P < 0.001$ | $F_{48, 576} = 9.889, P < 0.001$ |
| Fig.6E,<br>delta<br>T<sub>core</sub> | $F_{1, 12} = 3.580, P = 0.08$ | $F_{5.095, 61.14} = 46.72, P < 0.001$ | $F_{48, 576} = 10.66, P < 0.001$ |

A recent study established that capsaicin’s effects on body temperature are mediated by activation of spinal afferent pathways, rather than by direct modulation of peripheral targets such as smooth muscle cells (Yue *et al*., 2022). However, the central afferent CNS target that mediates robust capsaicin-induced hypothermia remain unknown.

The preoptic area POA is a central hub for thermoregulatory (Tan & Knight, 2018; Morrison & Nakamura, 2019; Rothhaas & Chung, 2021; Upton *et al*., 2021). Several studies have identified overlapping populations of predominantly Vglut2-positive (excitatory) neurons centred around the ventromedial preoptic area (VMPO), defined by distinct molecular markers, whose activation reliably induces profound hypothermia (Tan *et al*., 2016; Yu *et al*., 2016; Zhao *et al*., 2017; Hrvatin *et al*., 2020; Takahashi *et al*., 2020; Zhang *et al*., 2020). While the duration of hypothermia varied with the strength and duration of chemogenetic or optogenetic manipulations—and typically outlasted the hypothermic response induced by capsaicin—the depth and overall magnitude of the core and BAT temperature drop and tail vasodilation were intriguingly similar across these conditions.

We confirmed these earlier findings by chemogenetically activating VMPO^LepR^ neurons using a Gq-DREADD approach in LepR-Cre recombinase- expressing mice (Fig. 1C). CNO administration caused a pronounced decrease in core body temperature, from 35.69 ± 0.29 °C to 26.78 ± 0.31 °C at 1 h and to 25.15 ± 0.26 °C at 3 h, whereas saline injection had no effect (Fig. 1D, Table 1). This hypothermic response was accompanied by tail vasodilation and suppression of brown adipose tissue thermogenesis —similar to the responses evoked by capsaicin— as well as characteristic postural extension (Fig. 1E).

Together, these observations prompted the hypothesis that capsaicin-activated primary sensory afferents innervating cutaneous, muscular, and visceral tissues may engage LepR-expressing VMPO neurons to drive hypothermia. To test whether VMPO^LepR^ neurons contribute to the hypothermic effects of capsaicin, we bilaterally injected either a Cre-dependent AAV encoding tetanus toxin light chain (TeTxLC) to block synaptic transmission or a control virus into the POA of LepR-Cre mice (Fig. 2A, 2B). TeTxLC cleaves SNARE proteins, including vesicle-associated membrane protein 2 (VAMP2), thereby blocking synaptic neurotransmitter release (Sweeney *et al*., 1995; Wang *et al*., 2018).

**Figure 2.**
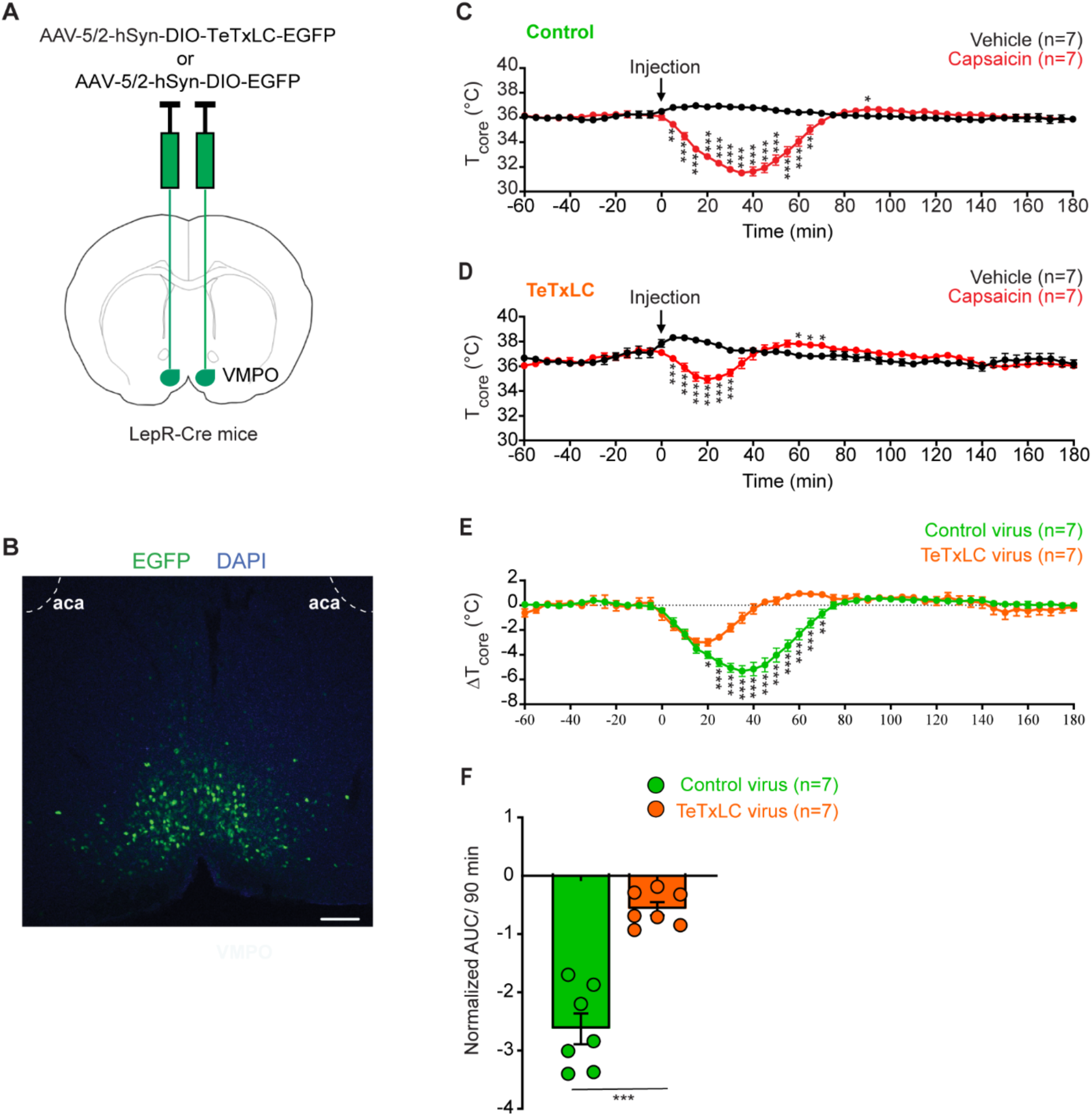
Silencing VMPO^LepR^ neurons partially attenuates capsaicin-induced hypothermia. **(A)** Schematic coronal diagram showing injection of AAV5/2-hSyn-DIO-TeTxLC-EGFP or AAV5/2-hSyn-DIO-EGFP into the POA of LepR-Cre mice. **(B)** Histological image showing TeTxLC-EGFP expression in LepR neurons in the VMPO area. Scale bar: 200 µm. **(C)** Core body temperature following a single subcutaneous injection of capsaicin (1 mg/kg) or vehicle in mice expressing EGFP in VMPO^LepR^ neurons (n = 7). **(D)** Core body temperature following a single subcutaneous injection of capsaicin (1 mg/kg) or vehicle in mice expressing TeTxLC in VMPO^LepR^ neurons (n = 7). **(E)** ΔTcore (capsaicin − vehicle) in control mice expressing EGFP and experimental mice expressing TeTxLC in VMPO^LepR^ neurons (calculated from data shown in panels C and D). Each data point corresponds to the mean value of body temperature ± SEM. Data were recorded every 5 min. Repeated-measures two-way ANOVA, followed by post-hoc Fisher’s LSD tests were performed. **(F)** Normalized area under the curve (AUC) of ΔTcore between 0 to 90 min in EGFP-expressing and TeTxLC-expressing mice in VMPO^LepR^ neurons. Statistical comparisons between two groups were performed using an unpaired two-tailed t-test with Welch’s correction. Asterisks indicate a statistically significant difference (*p < 0.05, **p < 0.01, ***p < 0.001).

Four weeks after AAV injection, we monitored core body temperature following subcutaneous administration of saline or capsaicin. As expected, capsaicin reduced core body temperature by 4.56 °C (from 36.07 ± 0.30 °C to 31.51 ± 0.24 °C) in control mice expressing EGFP in VMPO^LepR^ neurons (Fig. 2C, Table 1). In contrast, silencing VMPO^LepR^ neurons significantly attenuated, but did not abolish, the hypothermic response to capsaicin, resulting in a reduction of only 2.17 °C (from 37.11 ± 0.24 °C to 34.94 ± 0.30 °C) (Fig. 2D, Table 1). Analysis of the core body temperature difference (ΔT_core_ = T_core(capsaicin)_ - T_core(vehicle)_) revealed a pronounced hypothermic response in control mice expressing EGFP in VMPO^LepR^ neurons. In contrast, this hypothermic effect was markedly attenuated in mice with TeTxLC-mediated silencing of VMPO^LepR^ neurons, resulting in a significantly smaller area under the cure (AUC) (Fig. 2E, 2F, Table 1). These results indicate that VMPO^LepR^ neurons contribute to, but are not solely responsible for, the capsaicin-induced hypothermia effect.

### POA^Vglut2^ neurons are necessary for capsaicin-induced hypothermia

Previous studies suggest that hypothermia is primarily driven by Vglut2-positive preoptic neurons, rather than by Vgat-positive neurons (Song *et al*., 2016; Yu *et al*., 2016; Abbott & Saper, 2017; Upton *et al*., 2021). Because Vglut2 expression extends beyond LepR-positive neurons and is not restricted to the VMPO, we refer to this broader population as POA^Vglut2^ neurons. To test whether POA^Vglut2^ neurons mediate the hypothermic effect of capsaicin, we bilaterally injected either Cre-dependent AAV–TeTxLC or control virus into the POA of Vglut2-Cre mice to silence synaptic output (Fig. 3A, B).

**Figure 3.**
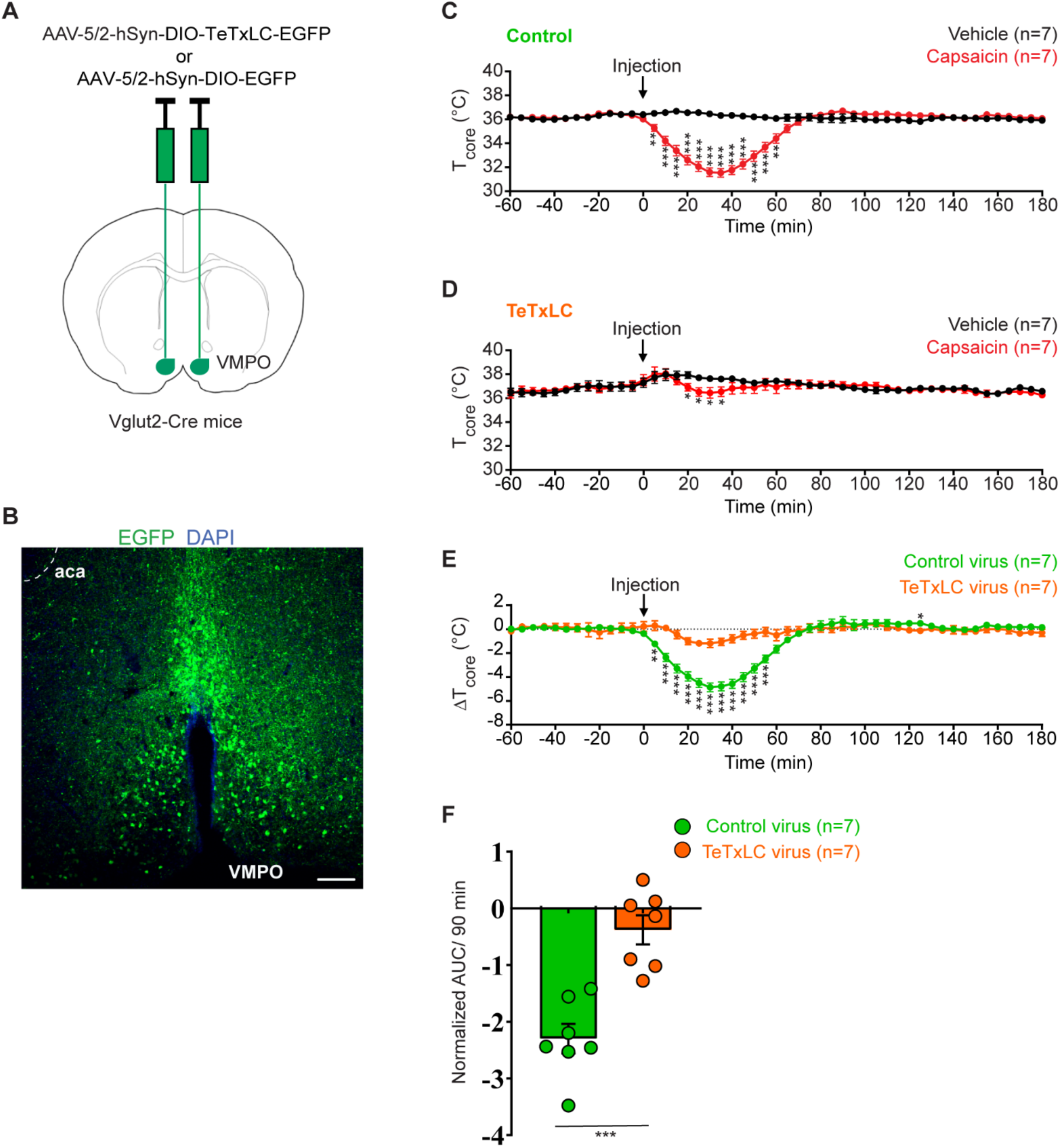
Silencing POA^Vglut2^ neurons effectively abolishes capsaicin-induced hypothermia. **(A)** Schematic coronal diagram showing injection of AAV5/2-hSyn-DIO-TeTxLC-EGFP or AAV5/2-hSyn-DIO-EGFP into the POA of Vglut2-Cre mice. **(B)** Histological image showing TeTxLC-EGFP expression in POA^Vglut2^ neurons. Scale bar, 200 µm. **(C)** Core body temperature following a single subcutaneous injection of capsaicin (1 mg/kg) or vehicle in mice expressing EGFP in POA^Vglut2^ neurons (n = 7). **(D)** Core body temperature following a single subcutaneous injection of capsaicin (1 mg/kg) or vehicle in mice expressing TeTxLC in POA^Vglut2^ neurons (n = 7). **(E)** ΔTcore (capsaicin − vehicle) in control mice expressing EGFP and experimental mice expressing TeTxLC in POA^Vglut2^ neurons (calculated from data shown in panels C and D). Each data point corresponds to the mean value of body temperature ± SEM. Data were recorded every 5 min. Repeated-measures two-way ANOVA followed by post-hoc Fisher’s LSD tests were performed. **(F)** Normalized area under the curve (AUC) of ΔTcore between 0–90 min in EGFP-expressing and TeTxLC-expressing mice in POA^Vglut2^ neurons. Statistical comparisons between two groups were performed using an unpaired two-tailed t test with Welch’s correction. Asterisks indicate statistically significant differences (*p < 0.05, **p < 0.01, ***p < 0.001).

In EGFP-expressing control mice, subcutaneous capsaicin induced a robust drop in core body temperature of 4.49 °C (from 36.02 ± 0.23 °C to 31.53 ± 0.36 °C) (Fig. 3C, Table 1). Strikingly, silencing POA^Vglut2^ neurons nearly abolished this hypothermic response, with core body temperature decreasing only minimally, from 37.51 ± 0.46 °C to 36.44 ± 0.44 °C (Fig. 3D, Table 1). AUC analysis of the change in core body temperature (capsaicin minus vehicle) revealed a robust hypothermic response in EGFP-expressing control mice. In contrast, silencing POA^Vglut2^ neurons with TeTxLC almost completely abolished this response and markedly reduced the AUC (Fig. 3E, F, Table 1).

These findings demonstrate that capsaicin-induced hypothermia is mediated largely via primary afferent pathways that activate Vglut2-positive POA neurons.

We noted that permanent TeTxLC-mediated silencing of POA^Vglut2^ neurons disrupted water balance and the mice exhibited pronounced polyuria and excessive drinking behavior (Fig. 4A, Table 1), a phenotype not observed following inhibition of VMPO^LepR^ neurons (data not shown). Food intake also showed a slight increase, but body weight was not significantly affected (Fig. 4B, C, Table 1).

**Figure 4.**
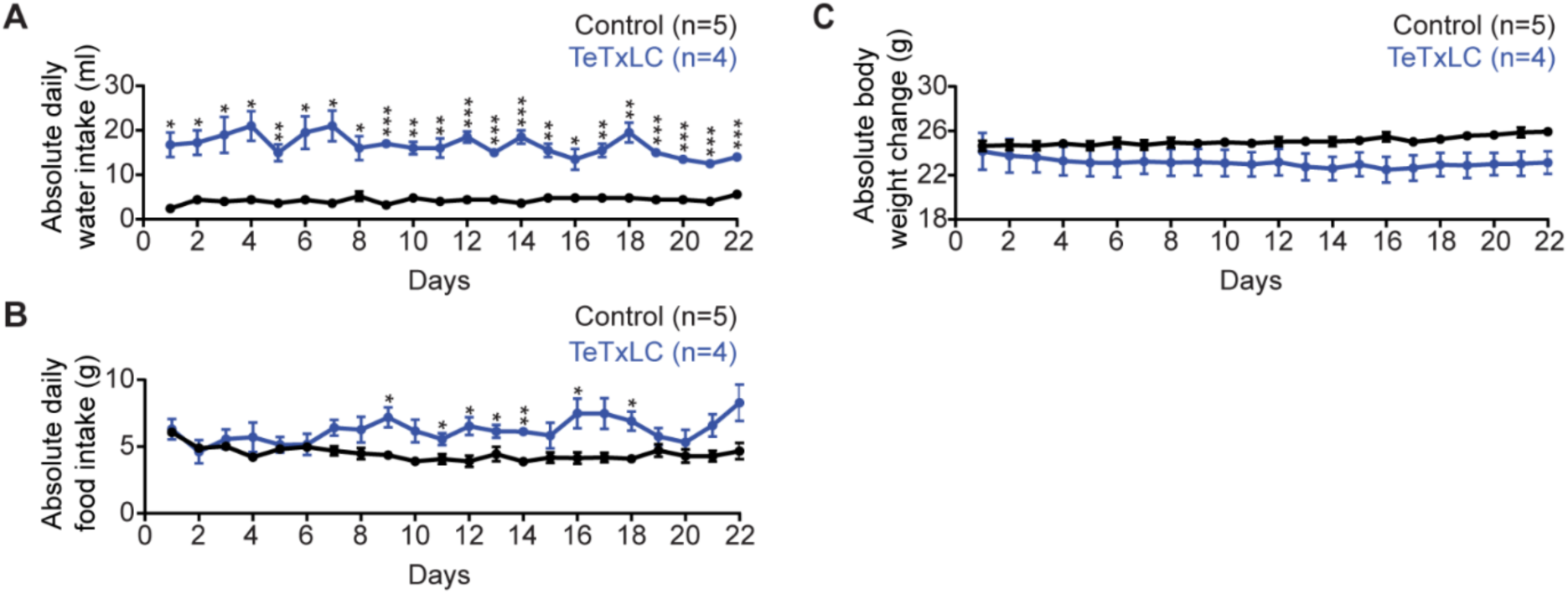
Silencing POA^Vglut2^ neurons increases water intake. **(A)** Daily water intake of mice expressing TeTxLC (n = 4) or expressing EGFP in POA^Vglut2^ neurons (n = 5). **(B)** Daily food intake of mice expressing TeTxLC (n = 4) or expressing EGFP in the POA^Vglut2^ neurons (n = 5). **(C)** Body weight of mice expressing TeTxLC or EGFP in POA^Vglut2^ neurons. All data are shown as the mean ± SEM. Repeated-measures two-way ANOVA, followed by post-hoc Fisheŕs LSD tests were performed. Asterisks indicate a statistically significant difference (\**p* < 0.05, \*\**p* < 0.01, \*\*\**p* < 0.001).

To avoid potential confounds from long-term silencing, we next employed a reversible chemogenetic approach. We stereotactically injected a Cre-dependent inhibitory DREADD (hM4Di–mCherry) into the POA of Vglut2-Cre mice (Fig. 5A, B), allowing transient suppression of neuronal activity (Sternson & Roth, 2014).

**Figure 5.**
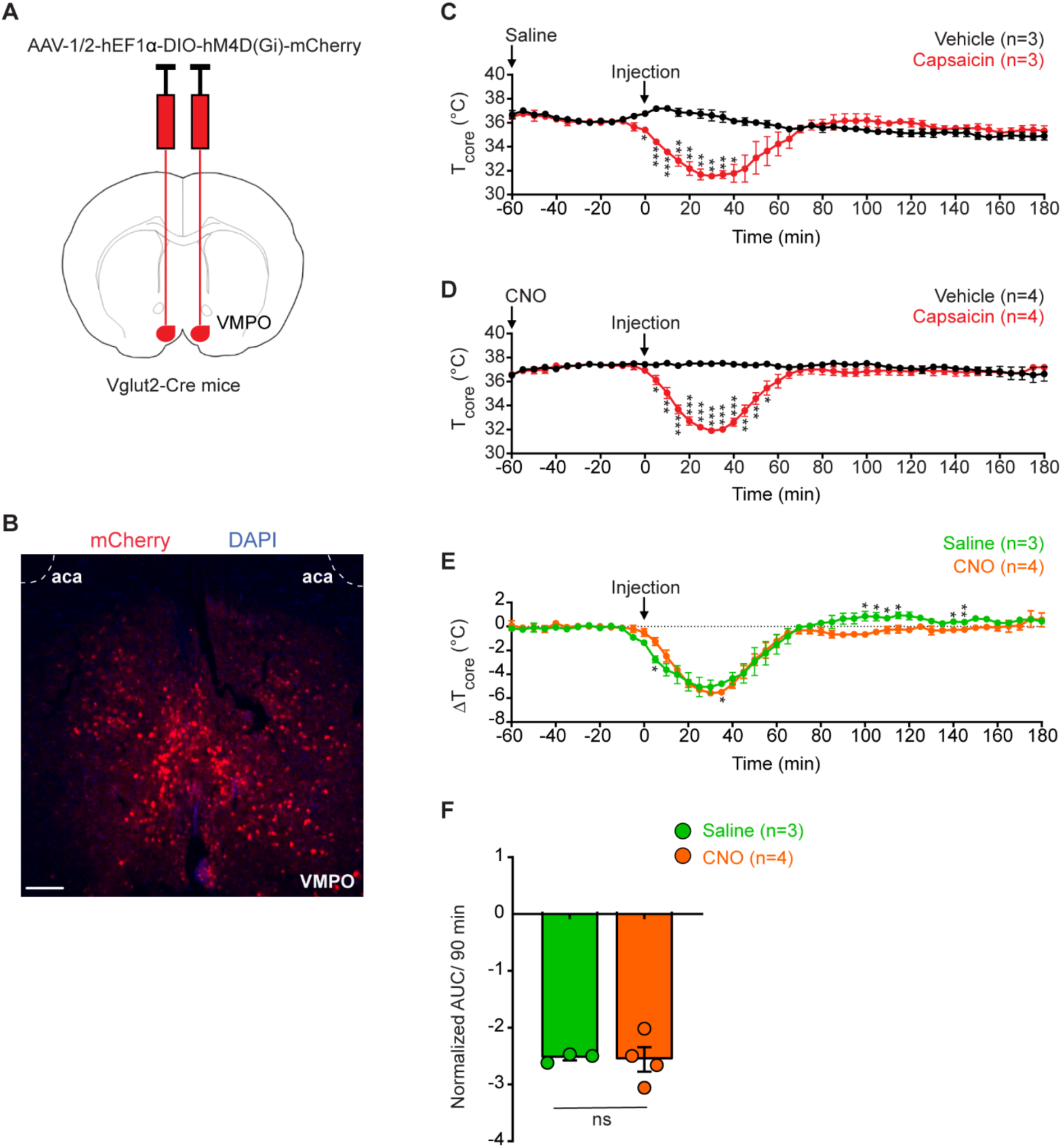
Chemogenetic inhibition of POAVglut2 neurons does not block capsaicin-induced hypothermia. **(A)** Schematic coronal brain section diagram showing injection of AAV-1/2–hEF1α-DIO-hM4D(Gi)-mCherry into the POA of Vglut2-Cre mice. **(B)** Histological image showing hM4D(Gi)-mCherry expression in Vglut2 neurons in the POA area. Scale bar: 200 μm. **(C)** Core body temperature following intraperitoneal saline injection (at t = -60 min) and subsequent subcutaneous administration of capsaicin (1 mg/kg) or vehicle (at t = 0 min) in mice expressing hM4D(Gi) in POAVglut2 neurons (n = 3). **(D)** Core body temperature following intraperitoneal CNO injection (at t = -60 min) and subsequent subcutaneous administration of capsaicin (1 mg/kg) or vehicle (at t = 0 min) in mice expressing hM4D(Gi) in POAVglut2 neurons (n = 4). **(E)** ΔTcore (capsaicin − vehicle) in saline- and CNO-treated mice. Each data point represents mean ± SEM (calculated from data shown in panels C and D). Data were recorded every 5 min. Repeated-measures twoway ANOVA, followed by post-hoc Fisheŕs LSD tests were performed. **(F)** Normalized area under the curve (AUC) of ΔTcore between 0–90 min for saline- and CNO-treated mice. Statistical comparisons between two groups were performed using an unpaired two-tailed t-test with Welch’s correction. Asterisks indicate a statistically significant difference (*p < 0.05, **p < 0.01, ***p < 0.001).

Mice pre-treated with saline showed a rapid hypothermic response to capsaicin, with core body temperature decreasing from 35.39 ± 0.04 °C to 31.54 ± 0.12 °C within 30 min (Fig. 5C, Table 1). Unexpectedly, CNO-mediated inhibition of POA^Vglut2^ neurons failed to block this response, as capsaicin still induced a pronounced decrease in core body temperature from 36.93 ± 0.21 °C to 31.90 ± 0.22 °C (Fig. 5D, Table 1). Capsaicin induced a strong hypothermic response in mice under both saline and CNO conditions, as reflected by the delta of core body temperature (capsaicin – vehicle). AUC analysis revealed no difference between treatments, indicating that CNO does not alter the capsaicin-induced hypothermia mediated by POA^Vglut2^ neurons (Fig. 5E, F, Table 1).

### POA^Vgat^ neurons partially mediate the hypothermic effect of capsaicin

To investigate whether POA^Vgat^ neurons contribute to capsaicin-induced hypothermia, we bilaterally injected either Cre-dependent AAV–tetanus toxin (TeTxLC) or control virus into the POA of Vgat-Cre mice (Fig. 6A, B). Four weeks after AAV injection, core body temperature was monitored following subcutaneous administration of saline or capsaicin.

**Figure 6.**
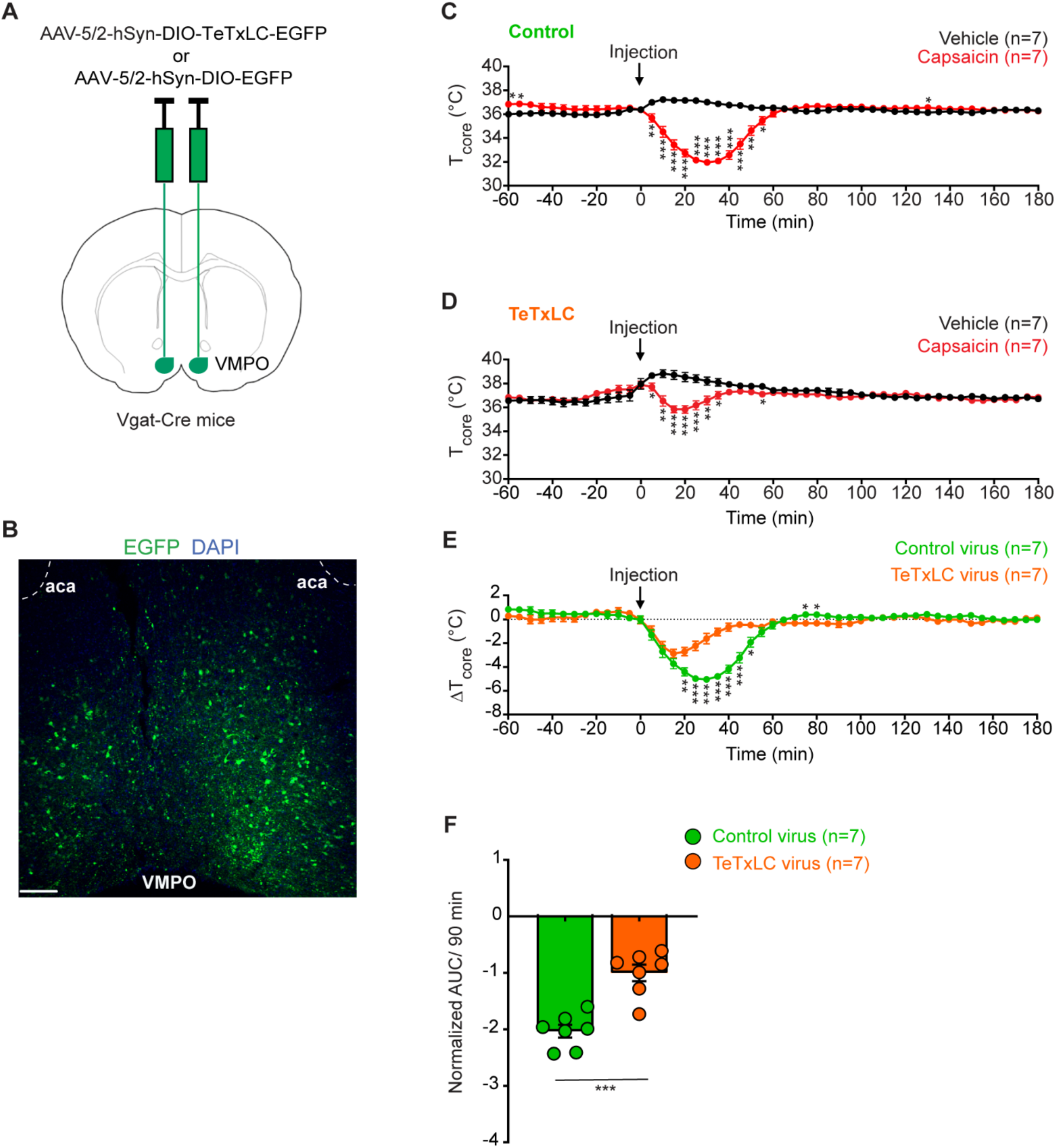
Silencing POA^Vgat^ neurons partially attenuates capsaicin-induced hypothermia. **(A)** Schematic coronal diagram showing injection of AAV5/2-hSyn-DIO-TeTxLC-EGFP or AAV5/2-hSyn-DIO-EGFP into the POA of Vgat-Cre mice. **(B)** Histological image showing TeTxLC-EGFP expression in POA^Vgat^ neurons. Scale bar, 200 µm. **(C)** Core body temperature following a single subcutaneous injection of capsaicin (1 mg/kg) or vehicle in mice expressing EGFP in POA^Vgat^ neurons (n = 7). **(D)** Core body temperature following a single subcutaneous injection of capsaicin (1 mg/kg) or vehicle in mice expressing TeTxLC in POA^Vgat^ neurons (n = 7). **(E)** ΔTcore (capsaicin − vehicle) in control mice expressing EGFP and experimental mice expressing TeTxLC in POA^Vgat^ neurons (calculated from data shown in panels C and D). Each data point corresponds to the mean value of body temperature ± SEM. Data were recorded every 5 min. Repeated-measures two-way ANOVA followed by post hoc Fisher’s LSD tests were performed. **(F)** Normalized area under the curve (AUC) of ΔTcore between 0–90 min in EGFP-expressing and TeTxLC-expressing mice in POA^Vgat^ neurons. Statistical comparisons between two groups were performed using an unpaired two-tailed t-test with Welch’s correction. Asterisks indicate statistically significant differences (*p < 0.05, **p < 0.01, ***p < 0.001).

In control mice expressing EGFP in POA^Vgat^ neurons, capsaicin induced a rapid and robust decrease in core body temperature within 30 min from 36.36 ± 0.19 °C to 31.96 ± 0.13 °C (Fig. 6C, Table 1). Silencing POA^Vgat^ neurons significantly attenuated, but did not abolish, the hypothermic response to capsaicin, resulting in a temperature decrease of 2.08 °C from 37.90 ± 0.30 °C to 35.82 ± 0.31 °C (Fig. 6D, Table 1). Calculation of the delta core body temperature (capsaicin – vehicle) revealed a pronounced hypothermic response in control mice expressing EGFP in POA^Vgat^ neurons. In mice with TeTxLC-mediated silencing of POA^Vgat^ neurons, this hypothermic response was reduced but not abolished, resulting in a smaller AUC compared with controls (Fig. 6E, F, Table 1). Different to blocking Vglut2-neuron output from the POA, silencing Vgat neuronal output did not result in abnormal water intake or secretion (data not shown). These results indicate that POA^Vgat^ neurons contribute partially to capsaicin-induced hypothermia, supporting the conclusion that peripheral TRPV1 activation engages multiple preoptic neuronal populations to mediate body cooling.

## Discussion

Our study demonstrates that capsaicin-induced hypothermia is mediated by afferent signals that converge onto neuronal circuits within the hypothalamic preoptic area (POA), a key integrative hub for thermoregulation. Rather than being driven by a single homogeneous neuronal population, our findings indicate that the hypothermic response emerges from circuit-level processing within the POA involving interacting excitatory and inhibitory neuronal components.

Since the first reports by Jancso-Gabor and colleagues (Jancso-Gabor *et al*., 1970a), a large body of literature reported that systemic application of capsaicin induces hypothermia (de Vries & Blumberg, 1989; Caterina *et al*., 2000; Christoph *et al*., 2008; Ayoub *et al*., 2011; Ivanova *et al*., 2023). However, the neuronal pathway(s) that mediate this response have remained elusive.

Several previous lines of evidence already implicated the hypothalamic preoptic area in the ensuing hypothermic response and found that capsaicin injections affected thermoregulatory responses of the POA. For example, POA warming, which triggers hypothermia similar to that of capsaicin injections was abolished or significantly decreased by prior capsaicin injections (Jancso-Gabor *et al*., 1970b; a). Additionally, after subcutaneous injection of capsaicin, the discharge frequency of the so-called warm-sensitive neurons in the POA increased (Nakayama *et al*., 1978).

Collectively, these and other studies (Sasamura *et al*., 1998; Karlsson *et al*., 2005) were taken as evidence that the POA participates in mediating capsaicin-induced hypothermia and that capsaicin’s site of action may be localized within the POA. However, intracerebral capsaicin injections ––including injections into the POA directly–– do not trigger pronounced hypothermia and are typically much less effective compared to peripheral administration routes such as oral gavage, subcutaneous, or intraperitoneal capsaicin injections (Jancso-Gabor *et al*., 1970b; de Vries & Blumberg, 1989; Caterina *et al*., 2000; Christoph *et al*., 2008; Ayoub *et al*., 2011; Inagaki *et al*., 2019; Ivanova *et al*., 2023). These early findings suggested that the primary site of action could be peripheral. Denervation of peripheral sensory neurons using resiniferatoxin, an ultra-potent capsaicin analog (Szallasi *et al*., 1989; Szallasi, 2023), or ablation of sensory neurons by genetic means (Mishra *et al*., 2011) prevented the capsaicin-induced hypothermia, also indicative of peripheral site of action. However, because these approaches could, in principle, also affect TRPV1-expressing neurons that may reside in the central nervous system through ablation or desensitization, they do not conclusively distinguish whether peripheral or central TRPV1 mediates body temperature control.

Additional pharmacological evidence for a peripheral site of action came from studies that employed peripherally restricted TRPV1 antagonists. Blocking TRPV1 activity results in the opposite effect compared to that of capsaicin and increases body temperature, revealing tonic activity of the receptor in the absence of an exogenous stimulus. TRPV1 antagonists chemically modified to remain primarily peripheral showed a similar propensity to trigger hyperthermia as antagonists that effectively cross the blood–brain barrier. (Gavva *et al*., 2007; Gavva, 2008; Tamayo *et al*., 2008).

Decisive genetic evidence for the predominant role of TRPV1 in peripheral sensory neurons came from a study utilizing conditional TRPV1 knock-out mice. These experiments showed that TRPV1 in sensory neurons (but not in smooth muscle cells) regulates body temperature bidirectionally when engaged by agonists or antagonists (Yue *et al*., 2022). Combining precise peripheral nerve transections with the application of selective TRPV1 antagonists corroborated these findings (Garami *et al*., 2023).

Additionally, available evidence suggests that the thermoregulatory effects of systemic TRPV1 activation are mediated primarily by somatosensory pathways (originating from dorsal root– and trigeminal ganglia neurons and their neuronal processes) rather than by vagal sensory neurons. Consistent with this, TRPV1-positive vagal afferents projecting to the nucleus tractus solitarius (NTS) —although themselves capable of inhibiting BAT thermogenesis— appear to be dispensable for the inhibition of BAT thermogenesis evoked by systemic TRPV1 agonists (Mohammed *et al*., 2018). Likewise, TRPV1 antagonist-induced hyperthermia is independent of the abdominal vagal nerve and instead depends on somatic afferents of the abdominal wall, which relay signals through the dorsolateral funiculus (DLF) to the LPB–raphe pathway controlling thermogenesis (Garami *et al*., 2023).

It has been speculated that TRPV1 activation could modulate body temperature by a purely peripheral mechanism: the release of vasoactive mediators from sensory nerve endings acting on smooth muscle and/or endothelial cells could, in principle, evoke vasodilation of blood vessels to dissipate body heat, independent of any central signalling. However, the two studies above show that this is an unlike scenario, but instead primary afferent sensory neuron pathways via spinal and supra-spinal sites relay TRPV1’s bidirectional thermoregulatory signals (Yue *et al*., 2022; Garami *et al*., 2023). This raises the important question where these TRPV1 signals are integrated to modulate downstream “effector” organs for body temperature regulation.

Our findings suggest that Vglut2-positive neurons located in the POA play a critical role in mediating capsaicin’s hypothermia-inducing effect.

The POA has long been known for its central role in body temperature regulation. Several studies have identified POA neurons, demarcated by the expression of disparate and partially overlapping genetic “markers”, that promote strong and long-lasting hypothermia in mice when activated optogenetically or chemogenetically (Yu *et al*., 2016; Zhao *et al*., 2017; Tan & Knight, 2018; Morrison & Nakamura, 2019; Hrvatin *et al*., 2020; Takahashi *et al*., 2020; Rothhaas & Chung, 2021; Upton *et al*., 2021).

The spinal–parabrachial pathway is considered the main route by which peripheral temperature information reaches the hypothalamic thermoregulatory centre (Geerling *et al*., 2016; Yang *et al*., 2020; Norris *et al*., 2021) and the VMPO —comprising the rostral parts of the median preoptic nucleus (MnPO), POA, and organum vasculosum of the lamina terminalis (OVLT)— has been proposed to serve as a principal entry point for afferent thermal signals regulating body temperature (Tan *et al*., 2016; Yu *et al*., 2016; Zhao *et al*., 2017; Tan & Knight, 2018; Morrison & Nakamura, 2019). Importantly, TRPV1 has been implicated not only in the detection of noxious heat and inflammatory stimuli, but also of more subtle warming stimuli (Yarmolinsky *et al*., 2016; Abd El Hay *et al*., 2025). It therefore appears conceptually consistent that systemic capsaicin —mimicking a warming stimulus and activating TRPV1-positive afferents— recruits POA/VMPO neurons that have previously been shown to become activated by warming (Yu *et al*., 2016; Ambroziak *et al*., 2025; Machado *et al*., 2025), thereby providing a plausible route by which capsaicin induces hypothermia.

While our findings support that the bulk of TRPV1 thermoregulatory signals are conveyed via the spinal-parabrachial–POA pathway, chemogenetic inhibition of POA^Vglut2^ neurons failed to block capsaicin-induced hypothermia (Fig. 5). Several explanations may account for this observation. One possibility is the existence of a parallel pathway that relays a fraction of TRPV1-dependent thermoregulatory signals to downstream brain regions independently of POA^Vglut2^ neurons. However, this scenario appears unlikely, as irreversible silencing of the same neuronal population using TeTxLC completely abolished capsaicin-induced hypothermia (Fig. 3).

A more plausible explanation is that chemogenetic inhibition via hM4Di was insufficient to counteract the strong activation of POA neurons triggered by systemic capsaicin. Activation of Gi-coupled signalling downstream of hM4Di receptors may reduce neuronal excitability but may not fully suppress the robust afferent drive reaching the POA.

An additional, not mutually exclusive possibility relates to circuit-level organization within the preoptic area. While our results identify excitatory POA neurons as principal mediators of capsaicin-induced hypothermia, the partial attenuation observed when silencing VMPO^LepR^ and POA^Vgat^ neurons suggests that multiple preoptic neuronal populations contribute to this response. Thus, afferent thermal signals may be integrated within local preoptic microcircuits composed of interacting excitatory and inhibitory neurons before being relayed to downstream thermoregulatory effectors. In support of this view, we previously obtained indirect evidence for local inhibitory microcircuits within POA that mediate heat defence (Kamm *et al*., 2021), and earlier pharmacological studies have also strongly implicated inhibitory POA neurons in thermoregulatory control (Morrison & Nakamura, 2019). In such a framework, hM4Di-mediated suppression of somatic action potential firing may not fully disrupt circuit-level signal propagation if thermosensory inputs percolate through local POA networks via dendritic or other non–action potential-dependent signalling mechanisms.

We found that TeTxLC-mediated POA^Vglut2^ neuron silencing promoted voracious water consumption (polydipsia) (Fig. 4A) and polyuria in mice (data not shown), whereas food intake and body weight were only slightly perturbed (Fig. 4B, C). It is known that MnPO neurons, which very likely are targeted by our viral approach, are important for drinking behavior and fluid control (Abbott *et al*., 2016; Abbott & Saper, 2017; Allen *et al*., 2017). Previous studies showed that acute optogenetic activation of glutamatergic MnPO neurons drives drinking behavior (Abbott *et al*., 2016; Abbott & Saper, 2017). In light of these findings, it is somewhat surprising that permanent silencing of POA^Vglut2^ neurons also promotes excessive drinking. One possible explanation is that these neurons normally exert a tonic influence on fluid-balance circuits. Chronic disruption of this control may destabilize downstream homeostatic mechanisms and shift osmolality set points or related renal parameters regulating fluid homeostasis. In such a framework, both acute activation and long-term silencing could lead to increased drinking, albeit through distinct circuit mechanisms.

In rodents, systemic capsaicin administration induces a dramatic drop in core body temperature accompanied by metabolic depression, features that resemble aspects of torpor —a naturally occurring physiological state characterized by a reversible reduction in body temperature, metabolism, and energy expenditure— although the capsaicin-induced state is markedly more transient. It is therefore intriguing that systemic capsaicin appears to activate POA neurons overlapping with neuronal populations previously shown to induce torpor-like hypothermia (Hrvatin *et al*., 2020; Takahashi *et al*., 2020; Zhang *et al*., 2020; Upton *et al*., 2021).

However, systemic TRPV1 activation likely exerts multiple organism-wide effects on cardiovascular and metabolic physiology, and reported effects of capsaicin (or dihydrocapsaicin) on blood pressure and heart rate have been inconsistent across studies (Chahl & Lynch, 1987; Hachiya *et al*., 2007; Ono *et al*., 2011; Mohammed *et al*., 2018; Shirani *et al*., 2021). Thus, while systemic capsaicin may engage components of the neural circuitry underlying torpor-like hypothermia, it likely does not fully recapitulate the coordinated physiological program of natural torpor. For example, hallmark cardiovascular adaptations of torpor, such as pronounced heart rate depression, may either not be triggered by TRPV1 activation or may be masked by the complex systemic actions of capsaicin.

Whether torpor (or a torpor-like hypothermic state) can be induced not only in small animals such as rodents but also in larger model organisms and humans, and whether peripheral sensory neuron pathways can be exploited to induce it, is currently unknown.

While medically-harnessed torpor or suspended animation states are far-flung possibilities (at least currently), a more concrete application of systemic capsaicin administration (or of its derivatives) in clinical settings is the induction of mild therapeutic hypothermia (Fosgerau *et al*., 2010). One main advantage over conventional methods (e.g. catheter-based body cooling) is that homeostatic (sympathetic) responses, which are triggered by physical body cooling and that counteract the beneficial effects in stroke or heart attack patients, are inhibited when capsaicin is used (or, for that matter, when the respective upstream VMPO neurons are activated).

In a globalized world, the dietary consumption of capsaicin-rich, spicy food is widespread. However, traditionally spicy cuisine has been associated with warmer climates (Gutierrez & Simon, 2016; Mozsik, 2016; Romanovsky, 2016). While there may be multiple reasons for this, it has been speculated that dietary capsaicin may help to cope with heat (Jancso-Gabor *et al*., 1970b; Szolcsanyi, 2015). It is therefore interesting to note that the POA neurons mediating capsaicin-induced body cooling have recently been shown by us and others to change plastically upon long-term heat exposure to drive heat acclimation and promote heat tolerance (Ambroziak *et al*., 2025; Cao *et al*., 2025).

It is therefore conceivable that frequent capsaicin application (or consumption) could have a similar ––albeit likely subtler–– effect, triggering POA neuron plasticity over longer periods, thereby aiding the body in adjusting to warm environments. Based on our study, future work may revisit the possibility that dietary capsaicin, or other pharmacologic routes that engage peripheral thermal pathways over extended periods, can be harnessed to foster beneficial acclimatory adaptation by driving hypothalamic plasticity.

## Acknowledgements

We thank Christine Siegmund, Amandine Cavaroc and Markus List for technical support; Dr. Katrin Schrenk-Siemens for inspiring discussions and critical reading of the manuscript; the Nikon Imaging Center at Heidelberg University for support with confocal microscopy. Funding: This work was supported by the European Research Council ERC-CoG-772395 and the German research Foundation SFB/TRR 152 and SFB1158.

## Author Contributions

J.S. and H.B. conceive the project. H.B. performed the experiments and data analysis. H.B. and J.S. wrote the manuscript.

## Methods

### Animals and housing conditions

Experiments were performed in accordance with the local ethics committee and approved by the local council (Regierungspräsidium Karlsruhe, Germany) under protocol number G-102/25. Mice were housed at an ambient temperature of 23 ± 1°C, relative humidity of 40-56%, and a 12-hour light/dark cycle (lights on at 7:00 a.m. and off at 7:00 p.m.). Throughout the housing and all experimental protocols, mice had *ad libitum* access to food and water.

Both male and female mice (8-12 weeks old) were used in the experiments. The mice lines used for the studies were B6N (C57BL/6N; Jackson Laboratory, IMSR_JAX 005304), LepRCre (B6.129-Lepr^tm3(cre)Mgmj/J^: Jackson Laboratory, IMSR_JAX 032457), Vglut2-ires-Cre (Slc17a6 ^tm2(cre)Lowl/J^; Jackson Laboratoray, IMSR_JAX 016963) and Vgat-ires-Cre (Slc32a1 ^tm2(cre)Lowl/J^; Jackson Laboratoray, IMSR_JAX 016962). Only heterozygous mice in the C57BL/6N background were used for the experiments.

### Stereotaxic injections of Adeno-Associated Virus (AAVs)

Stereotaxic surgeries were performed under aseptic conditions. Mice were anesthetized with Medetomidine (0.5 mg/kg), Midazolam (5 mg/kg), and Fentanyl (0.05 mg/kg) diluted in 0.9% sterile isotonic saline. Anesthetized animals were placed into a stereotaxic device (Model 1900, Kopf Instruments, USA) and kept on the heating pad set at 36°C. An ophthalmic ointment (Bepanthen; Bayer, Germany) was used to preserve corneal moisture. The head’s fur was removed using a depilatory cream, and the surgical site was cleaned and disinfected with a 7.5% povidone-iodine solution (Braunol; Braun, Germany). For local analgesia, 10% lidocaine (xylocaine, Pumpspray, Aspen Pharmacare) was applied. After exposing the skull bone via a small incision, bilateral burr holes were drilled through the skull with a hand drill (OS40, Osada Electric, Japan). A pulled glass micropipette with a 20-40 µm tip diameter was inserted into the brain. The virus was injected by a manual air pressure system. To reduce brain tissue damage, injections were performed slowly over 10 min period. For the POA, 250 nl of virus was injected in each side at the following coordinates from bregma AP: +0.800 mm, ML: ± 0.350 mm, DV: - 4.900 mm, and the glass micropipette was left in the site for an additional 5 min to permit diffusion into the parenchyma and then the micropipette was removed smoothly and slowly.

The scalp incision was closed using a 5-0 absorbable suture (MARLIN®, 17241041; Catgut, Germany) with a simple interrupted pattern. After the surgery, the anesthesia was antagonized using a subcutaneous injection of Antipamezol (2.5 mg/kg), Flumazenil (0.5 mg/kg), and Naloxon (1.2 mg/kg). For postoperative care, mice were injected subcutaneously with Carprofen at 5 mg/kg (Rimadyl; Zoetis, USA), kept on a heating pad at 37°C for 12 h in their home cage, and monitored closely to verify anesthesia recovery. Animals were allowed to recover from stereotaxic surgery for at least 4 weeks before experiments.

### Chemogenetic approach

For Designer Receptor Exclusively Activated by Designer Drug (DREADD) experiments, mice received an intraperitoneal injection of vehicle (saline) on day 1 and clozapine N-oxide (CNO; # C0832, Sigma-Aldrich) at the dose of 1.5 mg/kg on day 2. Of note, the injected volume of saline was similar to CNO to avoid any variation. Chemogenetic inhibition of POA^Vglut2^ neurons was carried out with vehicle (10% ethanol and 10% Tween-80 solution in saline) or capsaicin (10 mg/ml in vehicle) administration 60 minutes after saline or CNO treatment.

### Virus constructs

- ssAAV-DJ/2-hEF1α-dlox-hM3D(Gq)_mCherry(rev)-dlox-WPRE-hGHp(A) (Zurich Vector Core, 7.3 x 10E12 vg/ml).
- ssAAV-1/2-hEF1α-dlox-hM4D_mCherry(rev)-dlox-WPRE-hGHp(A) (Zurich Vector Core, 4.5 x 10E12 vg/ml).
- ssAAV-5/2-hSyn1-chI-dlox-EGFP_2A_FLAG_TeTxLC(rev)-dlox-WPRE-SV40p(A) (Zurich Vector Core, 7.7 x 10E12 vg/ml).
- ssAAV-5/2-hSyn1-dlox-EGFP(rev)-dlox-WPRE-hGHp(A) (Zurich Vector Core 1.1 x 10E13 vg/ml).

### Temperature transmitter implantation

Core body temperature was recorded using small and real-time readable temperature loggers (Anipill sysetem®; Bodycap, Paris, France). The logger weighed 1.7g and had a maximum diameter and height of 8.2 mm and 17.2 mm, respectively. Anipills had been programmed to record every 5 min. The intraperitoneal implantation of Anipill was performed under aseptic conditions and deep anesthesia. The fur of the abdomen area was shaved, the skin was disinfected with a 7.5% povidone-iodine solution (Braunol; Braun, Germany), and the cornea was protected with Bepanthen ointment (Bayer, Germany). A sterile Anipill logger was implanted in the abdominal cavity after an incision of 1 cm length was made in the skin and the abdominal muscles, using a subcutaneous injection of Antipamezol (2.5 mg/kg), Flumazenil (0.5 mg/kg), and Naloxon (1.2 mg/kg). For postoperative care, mice were injected subcutaneously with Carprofen at 5 mg/kg (Rimadyl; Zoetis, USA), and kept on a heating pad at 37°C for 12 h in their home cage and monitored closely to verify anesthesia recovery Thereafter, muscle layers and skin were sewed separately with 5-0 absorbable suture (MARLIN®, 17241041; Catgut, Germany). After the surgery, the anesthesia was antagonized. Animals were allowed to recover from surgery for at least 10 days before starting any further experiments.

### Capsaicin treatment

The stock solution was made by dissolving capsaicin ((*E*)-*N*-[(4-Hydroxy-3-methoxyphenyl) methyl]-8-methyl-6-nonenamide) (Tocris, # 0462) in 10% ethanol and 10% Tween-80 solution in saline at 10 mg/ml and then stored at -20°C. Immediately before the injection time, the stock solution was diluted with saline, and mice received a subcutaneous injection of capsaicin at a dose of 1 mg/kg or the vehicle solution in similar volumes.

### Immunohistochemistry

Mice were deeply anesthetized with isoflurane and then perfused intracardially with 20 ml phosphate buffer saline 0.1 M (PBS, pH 7.4) followed by 20 ml 4% paraformaldehyde (PFA). Brains were extracted and post-fixed overnight in 4% PFA and then cryoprotected with 30% sucrose in 0.1 M buffer phosphate for at least 3 days at 4°C. Brains were sectioned with a cryostat (CM3050, Leica, Germany) to obtain four series of 30 µm thick coronal sections at the level of the POA. The sections were collected in buffer phosphate saline (PBS-1X) with sodium azide (Sigma-Aldrich) and stored at 4°C. One series from each brain was used for EGFP and/or mCherry staining to verify the site of AAV microinjections, and the other 3 series were stored in an anti-freeze solution (Cryoprotectant, DEPC-treated, Bioenno Lifesciences).

For immunohistochemistry, free-floating brain sections were washed 3 times for 10 min in PBS-1X, pH 7.4, and incubated in 10% normal horse serum (Linaris Biologische Produkte GmbH) in PBS with 0.2% Triton X-100 (Merck) for 2 h. Primary antibody was diluted in 1% normal horse serum and 0.2% Triton X-100 in PBS, and sections were incubated for 24 h at 4°C under slow shaking. The following antibodies were used: Chicken anti-GFP (1:1000; Rockland, cat # 600-901-215S) and rabbit anti-mCherry (1:1000; Invitrogen, cat # PA5-34974). Afterward, floating sections were washed 3 times for 10 min in PBS-1X. Sections were then incubated with Alexa 488 anti-chicken (1:500, Thermo Fischer, cat # A-11039) and Alexa 555 anti-rabbit (1: 500, Thermo Fischer, cat # A-21430) for 2 h at room temperature. Finally, brain sections were rinsed 3 times for 10 min in PBS-1X, mounted onto glass microscope slides, and coverslipped with an Antifading Mounting Medium DAPI (ImmunoSelect®, Biozol Diagnostica Vertrieb GmbH). Fluorescent images were acquired at the Nikon imaging center at Heidelberg University, using the Nikon A1R confocal microscope under Nikon Plan Apo λ 10x magnification NA 0.45 (working distance 4 mm, the field of view 1.27 x 1.27 mm) objective.

### Statistical analysis

Data are presented as mean ± standard error of the mean (SEM). We assumed statistical significance at p < 0.05. No animals or data points were excluded, and each experiment was repeated 2 to 3 times. Statistical analysis was performed using GraphPad Prism 8 software (GraphPad Software, San Diego, CA). Two-way repeated measures analysis of variance (rm-ANOVA) followed by Fisher’s LSD *post hoc* test was applied to assess the effect of treatment (capsaicin vs vehicle), time, and the interaction between these factors.

